## Supplementary Data for "Cell fixation improves performance of *in situ* crosslinking mass spectrometry while preserving cellular ultrastructure"

Dr. David Schriemer

Department of Biochemistry and Molecular Biology, Cumming School of Medicine

University of Calgary, 3330 Hospital Drive NW T2N-4N1

---

[#]Both authors contributed equally to the work.

**Supplementary Information**

#### Contents

**Supplementary Figure 1.** Visualization of A549 cells during *in situ* crosslinking

**Supplementary Figure 2.** NHS-ester labeling in formaldehyde-fixed cells

**Supplementary Figure 3.** Method for increasing labeling yield

**Supplementary Figure 4.** CSM score distributions

**Supplementary Figure 5.** Predicted AlphaFold multimer structure of API5-ACIN1 interaction

**Table S1.** *In situ* crosslink breakdown for DSS and PhoX at 5% and 1% FDR

**Table S2.** Fixed 3X DSS 5% FDR PPI with protein information

**Table S3.** Fixed Perm PhoX 5% FDR PPI with protein information

### Supplementary Information

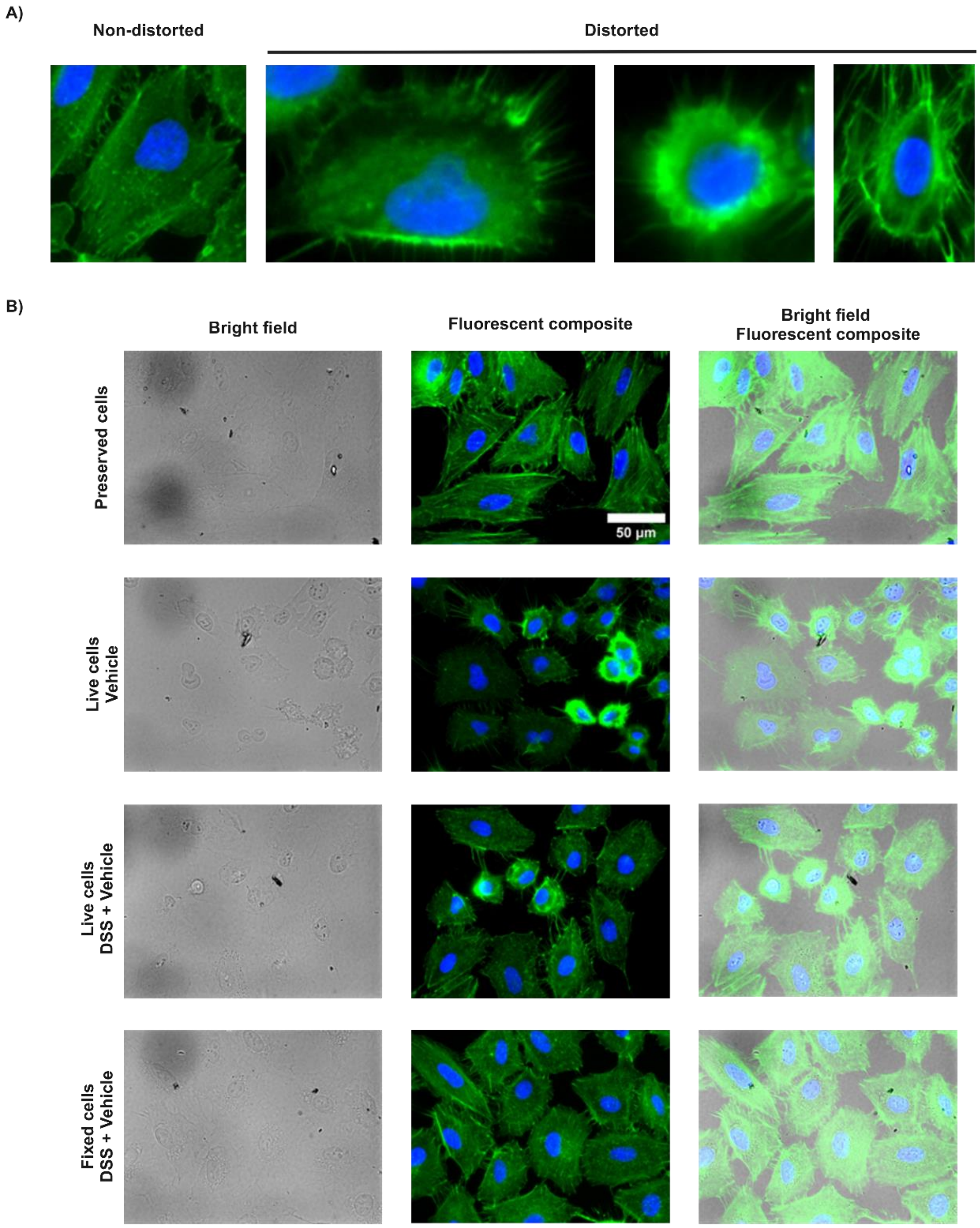


**Supplementary Figure 1. Visualization of A549 cells during *in situ* crosslinking.** (A) Representative examples of cells for the quantitation of cellular ultrastructure preservation. (B) Bright field and fluorescent images of formaldehyde-preserved cells, live cells treated with DMSO, live cells treated with DSS + DMSO, and formaldehyde-preserved cells treated with DSS + DMSO.

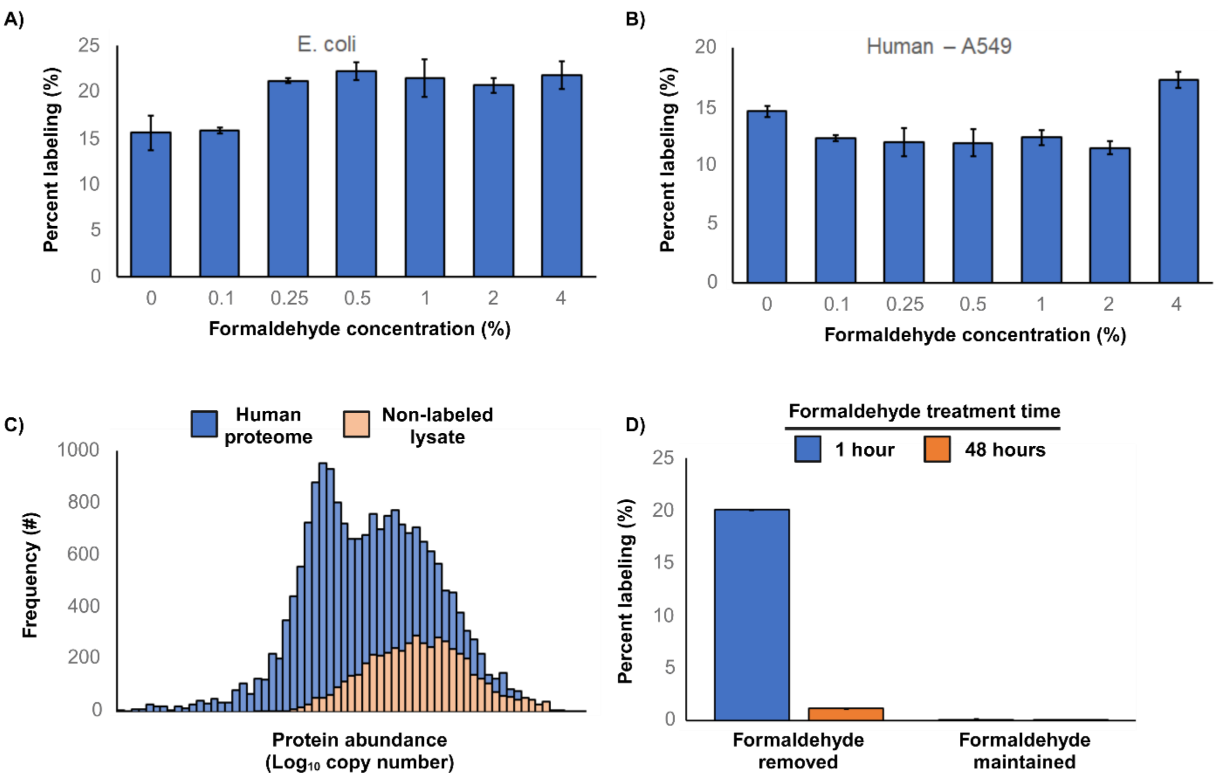


**Supplementary Figure 2. NHS-ester labeling in formaldehyde-fixed cells.** Labeling analysis of N-(propionyloxy)succinimide reactions with (A) *E. coli* and (B) human cells, over increasing concentrations of formaldehyde. (C) Protein abundance histogram of human proteome (blue) and protein ID's from MS-acquired non-labeled lysate (orange). Protein abundances retrieved from PaxDb[46]. (D) Labeling of N-(propionyloxy)succinimide in formaldehyde-fixed A549 cells with excess formaldehyde washed away or maintained, after1 hour (blue) or 48 hours (orange) fixation times.

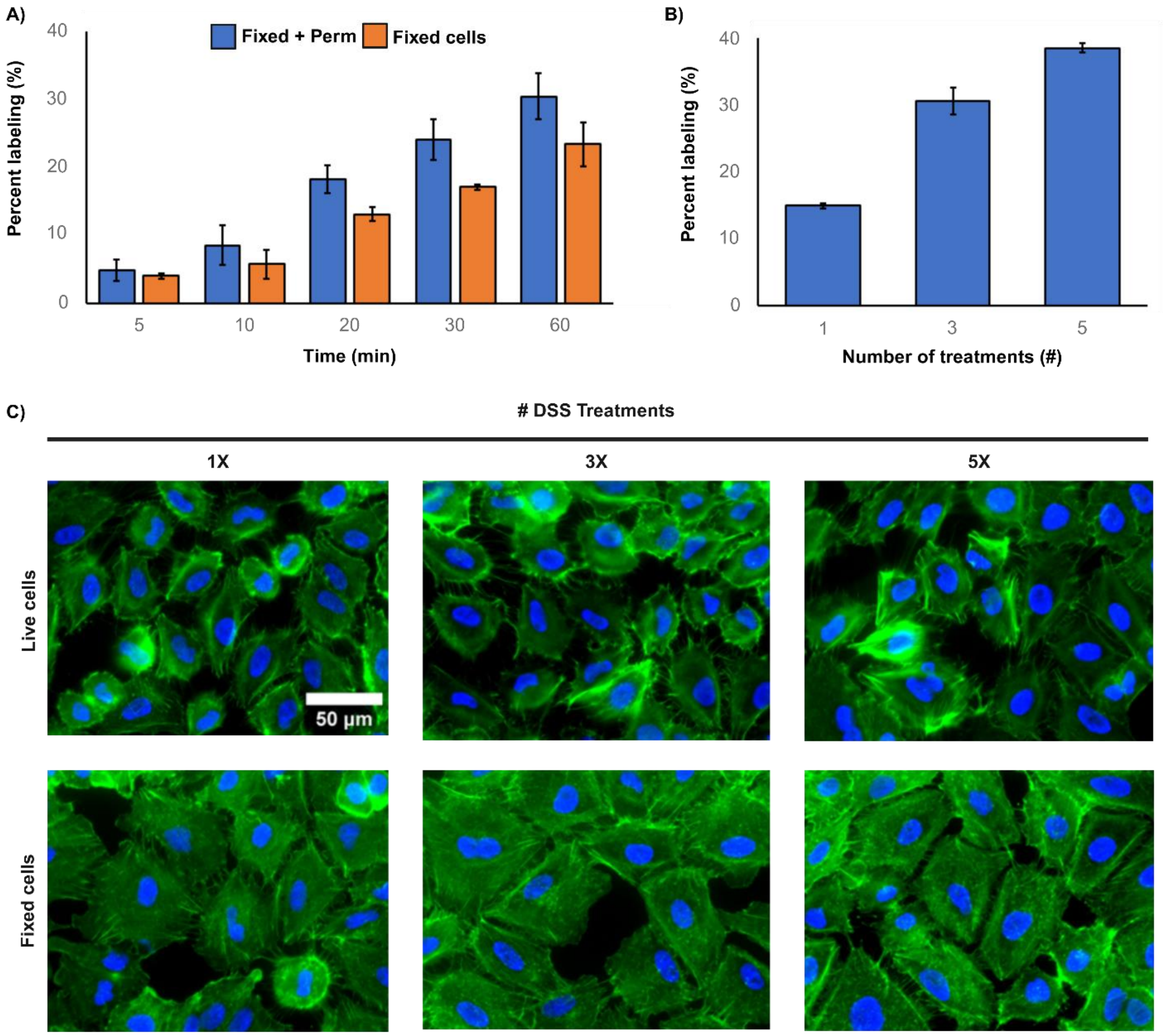


**Supplementary Figure 3. Method for increasing labeling yield.** (A) Time-course of biotin-X-NHS labeling in A549 cells that have been fixed (orange) or fixed and permeabilized (blue). (B) Percent labeling of increasing number of 1 mM treatments of biotin-X-NHS in fixed A549 cells. (C) Fluorescent micrographs of live and fixed A549 cells undergoing increasing numbers of DSS treatments, visualizing actin (green) and DNA (blue).

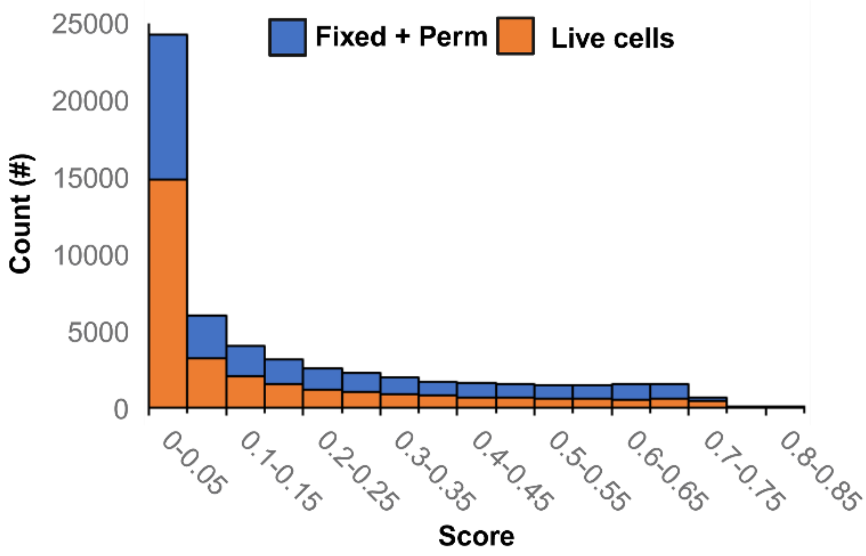


**Figure S4. CSM score distributions.** CSM score distributions of fixed + permeabilized (blue) and Live (orange) DSS treated A549 cells at 5% FDR. Scores obtained from pLink2 data output.

### Supplementary Information

**Model 1**

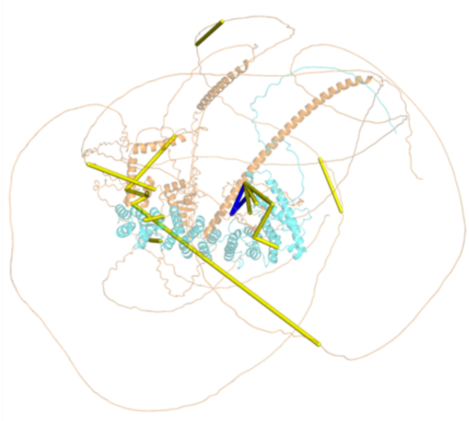

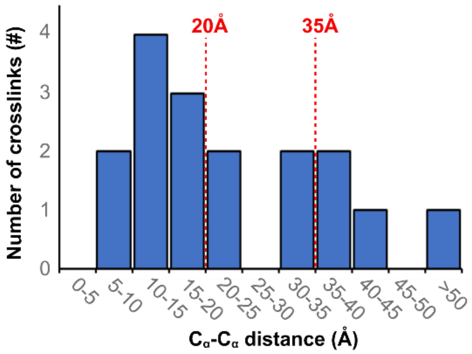


**Model 2**

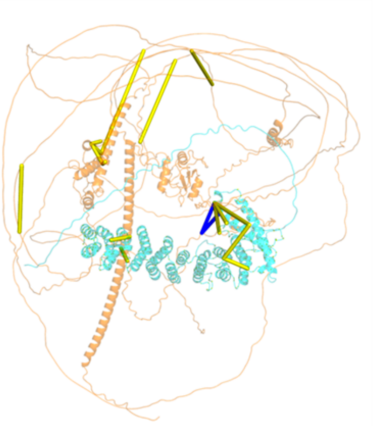

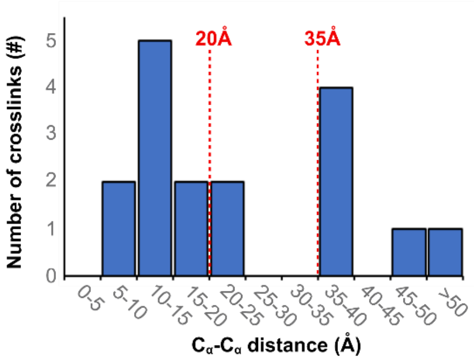


**Model 3**

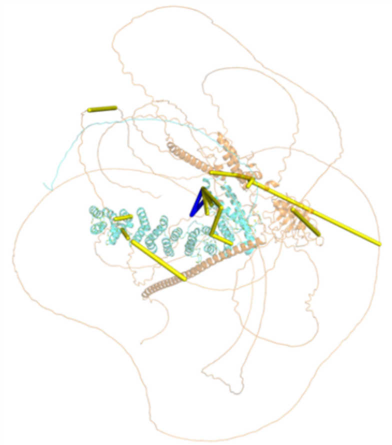

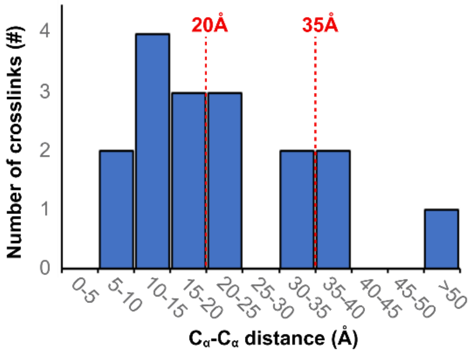

### Supplementary Information

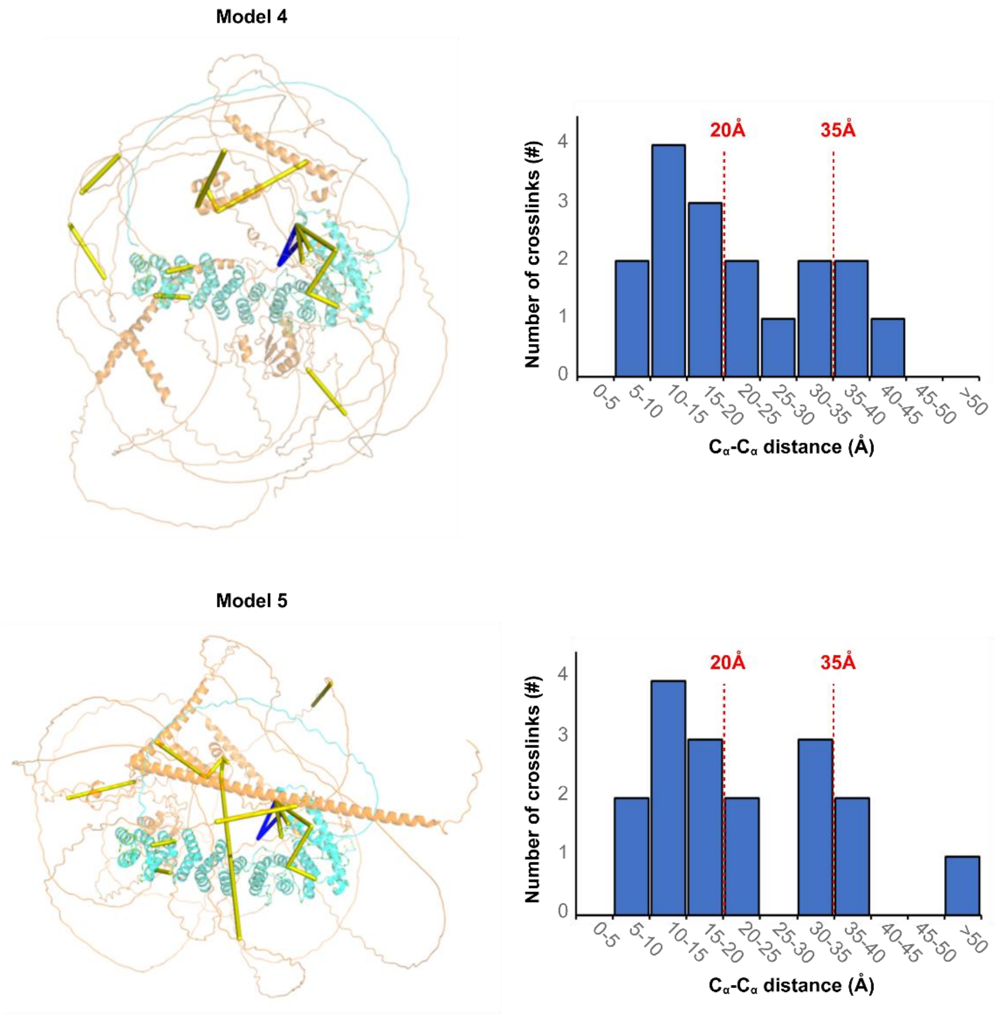


**Figure S5. Predicted AlphaFold multimer structure of API5-ACIN1 interaction.** Crosslinks mapped to 5 predicted models (left) and the corresponding $C_\alpha$-$C_\alpha$ distances histograms (right) of mapped PhoX crosslinks. API5 and ACIN1 coloured cyan and orange, respectively. Intra-protein and inter-protein crosslinks coloured yellow and blue, respectively.
